# Modeling cryo-EM structures in alternative states with generative AI and density-guided simulations

**DOI:** 10.1101/2025.02.06.636862

**Authors:** Tatiana Shugaeva, Rebecca J Howard, Nandan Haloi, Erik Lindahl

## Abstract

Modeling atomic coordinates into a target cryo-electron microscopy map is a crucial step in structure determination. Despite recent advances, proteins with multiple functional states remain a challenge - particularly when suitable molecular templates are unavailable for certain states, and the map resolution is not high enough to build de novo models. This is a common scenario, for example, among pharmacologically relevant membrane-bound receptors and transporters. Here, we introduce a refinement approach in which *i*) several initial models are generated by stochastic subsampling of the multiple sequence alignment (MSA) space in AlphaFold2, *ii*) the resulting models are subjected to structure-based clustering, *iii*) density-guided molecular dynamics simulations are performed from the centroid structures, and *iv*) a final model is selected on the basis of both map fit and model quality. This results in improved fitting accuracy compared to single starting point scenarios for three membrane proteins (the calcitonin receptor-like receptor, L-type amino acid transporter and alanine-serine-cysteine transporter which undergo substantial conformational transitions between functional states. Our results indicate that ensemble construction using generative AI combined with simulation-based refinement facilitates building of alternative states in several families of membrane proteins.

## Introduction

Cryo-electron microscopy (cryo-EM) is one of the most widely used and rapidly developing techniques for biomolecular structure determination. Advances in both hardware and algorithms have enabled density map reconstruction at or near atomistic resolution for numerous systems [1, 2]. In parallel, automated as well as manual model building methods have provided detailed insights into molecular processes, complex architectures and new conformations [3]. Despite this progress, building structures remains challenging for systems where 1) a target protein undergoes conformational transitions between multiple states [4] and 2) the map resolution is not high enough for de novo model building [5–8].

Density-guided molecular dynamics (MD) simulations, where a biasing potential is added to the classical forcefield to move atoms toward the experimental map, might provide solutions to both aforementioned challenges [9–14]. However, the success of density-guided MD simulation depends on the reliability of the initial model. Even when a template structure resolved in one state is available, it may differ substantially from a target density resolved in another state, such that automated approaches produce poorly fitting or nonphysical models (Fig. 1 - left). Such scenarios may be common, for example, among membrane-bound receptor and transporter proteins, which constitute overrepresented but dynamically complex pharmaceutical targets [15]. Accuracy may be further improved by fitting a well-sampled ensemble rather than a single initial model (Fig. 1 - right); however, the generation of such ensembles can be computationally expensive and time-consuming using classical MD or enhanced sampling simulations [16].

**Fig. 1.**
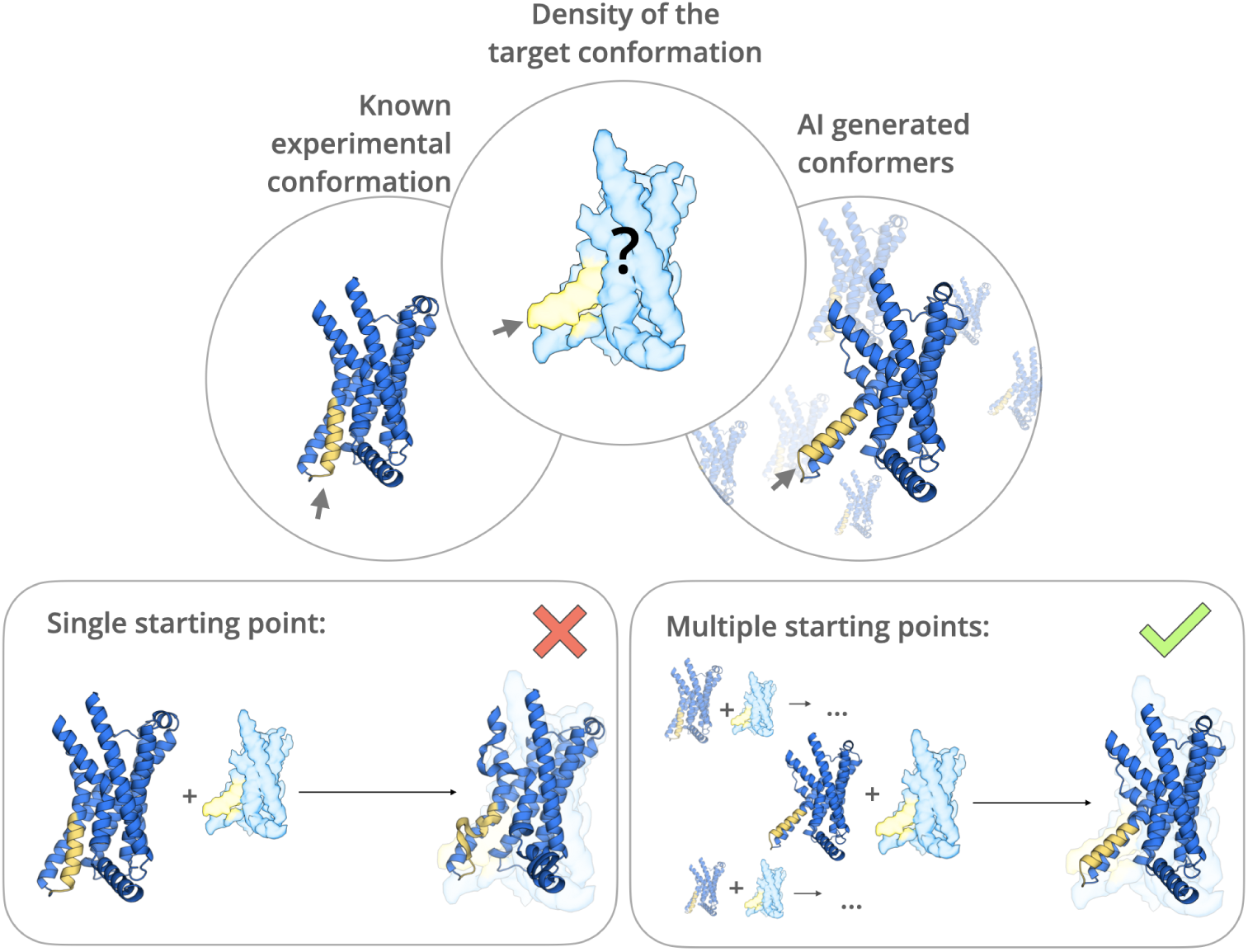
Challenges and strategies in automated modeling of proteins occupying alternative conformations in cryo-EM maps. *Left,* fitting of an experimental target density (surface) based on a previously known structure (ribbons) may be challenging for a given protein (blue) if its conformation differs in one or more regions (yellow). *Right,* generative AI methods enable simulation seeding from multiple prospective models (ribbons), one or more of which may more accurately approximate the target density (surface), and result in successful automated fitting.

Here, we present an approach (Fig. 2) that combines generative artificial intelligence (AI) methods and flexible fitting to refine protein structures based on sequence information and low-to medium-resolution cryo-EM data. We demonstrate the applicability to three test cases, all membrane proteins that undergo conformational change between experimentally resolved states. Taking advantage of recent developments in neural network-based structure prediction [17], we use stochastic subsampling of the multiple sequence alignment (MSA) depth in AlphaFold2 [18] to generate an ensemble of potential starting structures, then run density-guided MD simulations from cluster centroids to select an optimized structure for a given experimental density. We successfully resolve state-dependent differences including the bending of an individual helix, rearrangement of neighboring helices, and reformation of an entire domain, demonstrating the applicability to a range of systems and conformational dynamics.

**Fig. 2.**
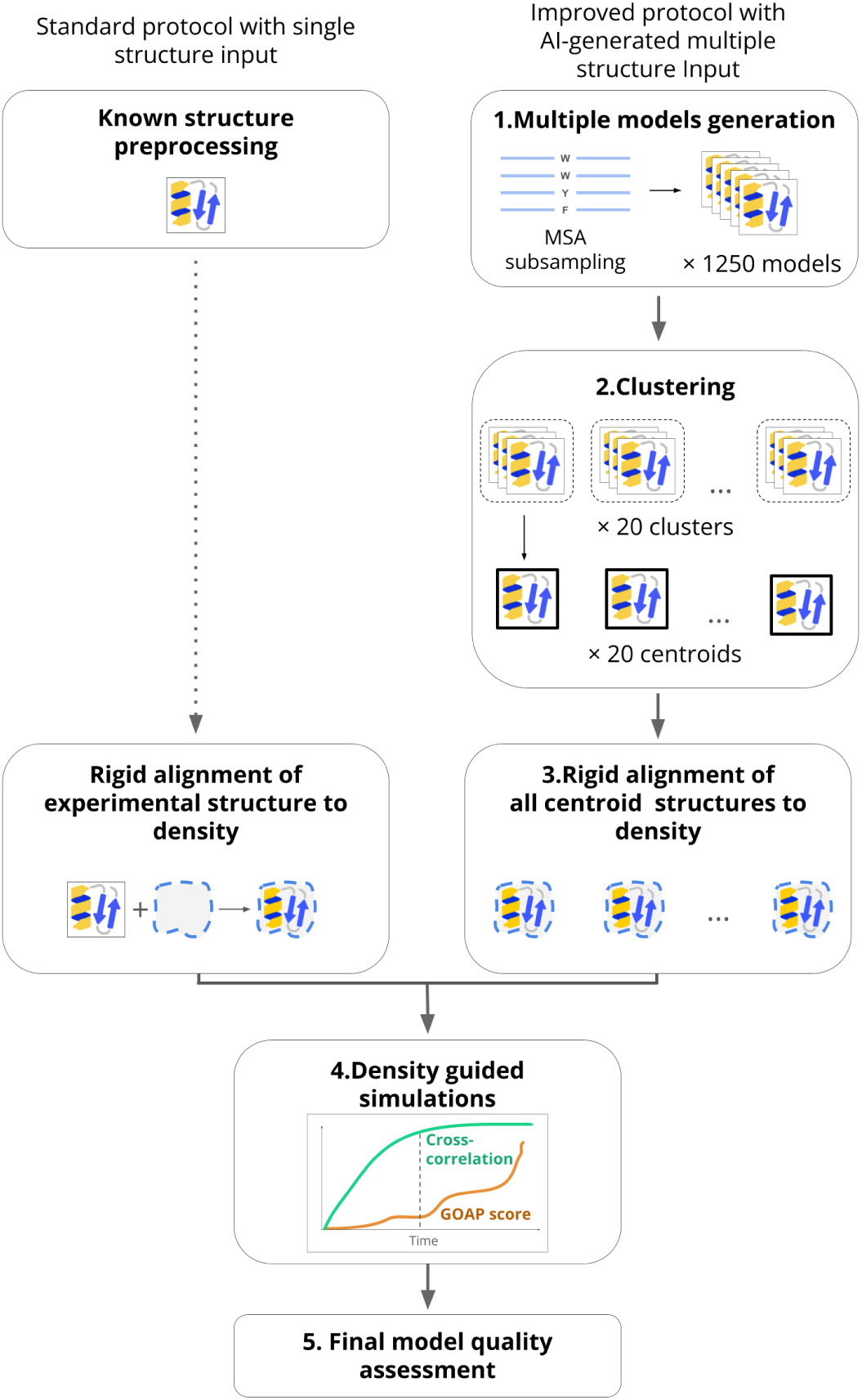
Incorporating generative AI models with flexible fitting. *Left,* in a standard approach to automated fitting of a new cryo-EM map, a known experimental structure is pre-processed, aligned to the target density, and subjected to density-guided MD simulations. *Right,* in the approach described here, an ensemble of models is generated by MSA subsampling in AlphaFold2 and clustered. Cluster centroids are aligned to the density and subjected to similar simulations. From the simulation with the highest mean cross-correlation to the target density, a final model is chosen to optimize map fit (cross-correlation) and model geometric quality (GOAP score). The protein emoji used in the scheme is taken from https://github.com/whitead/protein-emoji

## Results

### Incorporating generative models with flexible fitting

As a straightforward approach to incorporating generative AI models with flexible fitting in determining challenging protein structures (Fig. 2 - right), we first used stochastic subsampling of input MSA depth in AlphaFold2 to produce 1250 models for each of three protein test systems. To filter out substantially misfolded models, we prioritized those scoring better than (below) -100 on the basis of generalized orientation-dependent all-atom potential (GOAP) structure quality scoring (Methods). We then clustered these to identify a limited set of plausible models representative of the generated ensemble, and used the model closest to the cluster centroid for fitting to an experimental density, termed the target state. After rigid-body alignment to the relevant density, we subjected each centroid model to density-guided MD simulations, and selected the simulation with the highest mean cross-correlation to the target density. Across this centroid simulation, we evaluated the cross-correlation to the experimental map as a model fitting metric, and GOAP score as a structure geometry quality metric. After normalizing each metric to [0, 1], we subtracted the GOAP score from the cross-correlation to calculate a compound score for each simulation frame, with higher scores representing a combination of good fit (high correlation) and good geometric quality (minimal penalty e.g. due to overfitting causing clashes). We then selected the frame with the highest compound score as a final model.

To compare our approach to a more standard flexible fitting protocol [14], we performed parallel density-guided MD simulations for each experimental map using a single experimental structure of the same protein resolved under different conditions, termed the known state (Fig. 2 - left). This scenario is common in investigation of challenging targets, in which a structure has been determined in one functional state, but deviates substantially from an experimental density representing a different state. As for many experimental structures, for two of our three test systems fitting required interpolation of loops not reported in the initial known model, a step rendered unnecessary in our generated ensemble approach described above. In both ensemble-based and single-model approaches, we refrained from applying secondary structure restraints. Such restraints are commonly used in density-guided simulations to prevent nonphysical structure disruptions [10, 19], but can limit the accommodation of conformational transitions — for example, kinking of a helix, rearrangement among secondary structure elements, or remodeling of an entire domain.

We tested our approach by modeling publicly available experimental densities for three membrane proteins: the calcitonin receptor–like receptor (CLR, EMD 20906, 2.3 Å resolution) [20], L-type amino acid transporter (LAT1, EMD 30841, 3.4 Å resolution) [21] and alanine-serine-cysteine transporter 2 (ASCT2, EMD 12142, 3.4 Å resolution) [22] (Table 1). These structures were not present in the AlphaFold2 training set. To explore the applicability of our approach to medium-resolution maps, we added a 1 Å Gaussian blur to each target density. In each case, an initial known-state model for standard flexible fitting was chosen from an experimental structure determined in a different state than the target density (CLR, PDB ID 7KNT [23]; LAT1, PDB ID 6IRS [24]; ASCT2, PDB ID 6RVX [25]). For CLR, the known-state structure differed from the target-state map in the bending of a single helix; for LAT1, it differed in the arrangement of two neighboring helices; for ASCT2, it exhibited a substantial conformational transition involving most of the transmembrane helices (Fig. 3 - above). Subsequent to density-guided simulations, we used the deposited structure associated with each target state (CLR, PDB ID 6UVA [20]; LAT1, PDB ID 7DSQ [21]; ASCT2, PDB ID 7BCQ [22]) as a reference for fitting quality (Fig. 3 - below). Although such a structure would not be available in an anticipated application of our approach to a newly reconstructed density, it served as a valuable ground truth for validation of this work. We did not use the RMSD values as a criterion to select the final model from simulations.

**Fig. 3.**
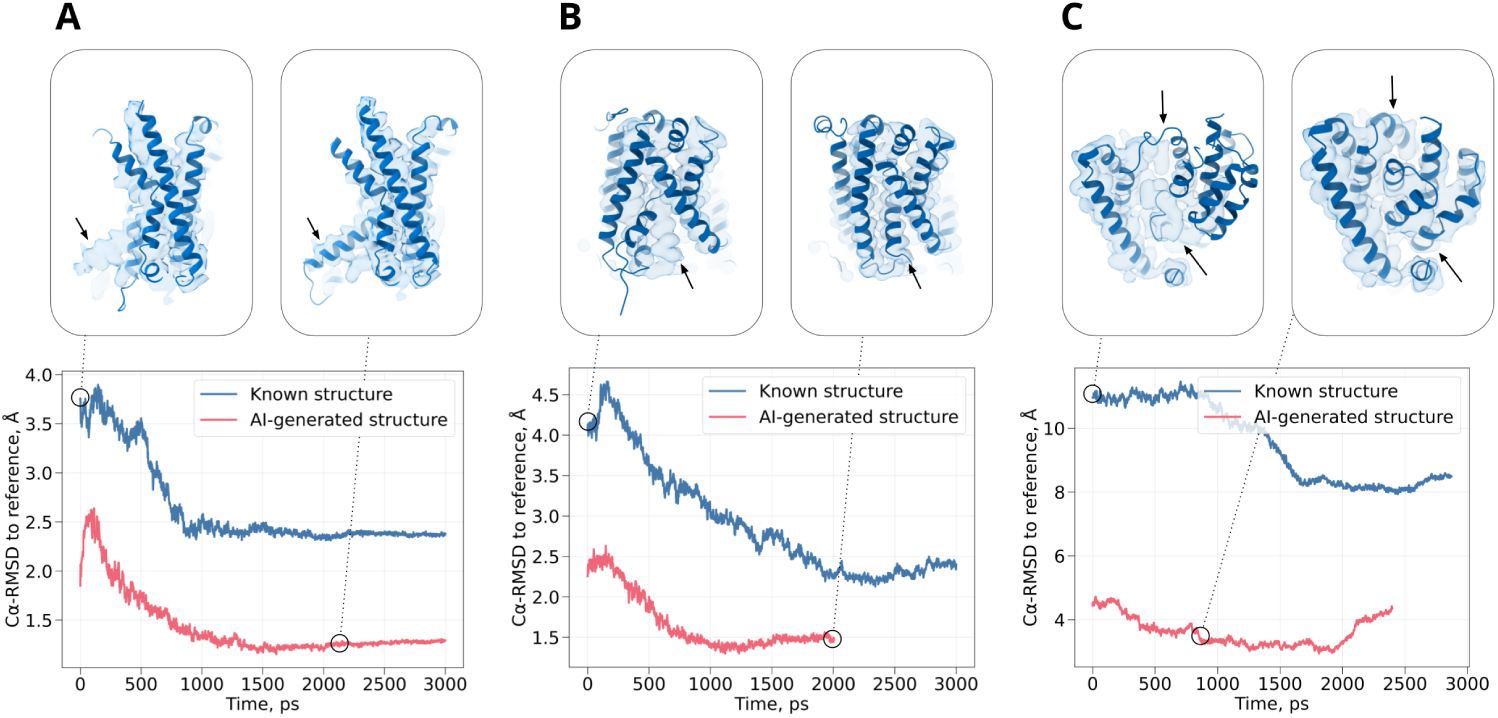
Refinement of an alternative protein state by AI model generation in three test systems: (A) CLR, (B) LAT1, (C) ASCT2. Plots track C*α* RMSD to the target structure during density-guided simulations starting either from the known structure (blue) or the best-fit cluster centroid from our generated ensemble (red). Upper insets show the known structure (*left* ) and final model from our approach (*right* ) for each system, aligned to the corresponding density. The arrow highlights the misaligned portion of the protein in the known structure.

**Table 1.** PDB and EMDB IDs for the systems used in the study.

|  | EMDB ID (target state) | PDB ID (target state) | PDB ID (known state) |
| --- | --- | --- | --- |
| CLR | 20906 | 6UVA | 7KNT |
| LAT1 | 30841 | 7DSQ | 6IRS |
| ASCT2 | 12142 | 7BCQ | 6RVX |

### Test case 1: helix bending in a 7-TM receptor

We first investigated the characteristic conformational transition of the G-protein coupled calcitonin receptor-like receptor (CLR), a component of both the calcitonin gene-related peptide receptor (CGRPR) and adrenomedullin (AM) receptor, involved in wound healing, vasodilation and other widespread physiological functions [26, 27]. A notable conformational transition between functional states of CLR transmembrane domain occurs in the 6th transmembrane helix (TM6, residues 329-354 in the target structure), which is relatively straight in the inactive state without G protein (Fig. 4A - left), but bends upon activation (Fig. 4A - right) [28]. This rearrangement remodels the intracellular binding interface at the N-terminal end of TM6, allowing it to interact with the Gs-protein *α* subunit. We used an inactive structure of CLR transmembrane domain, extracted from the larger CGRPR (PDB ID 7KNT)[23], as the known state. The cryo-EM density from an active state of CLR, extracted from the larger AM receptor (EMD 20906)[20], served as a target for model refinement.

**Fig. 4.**
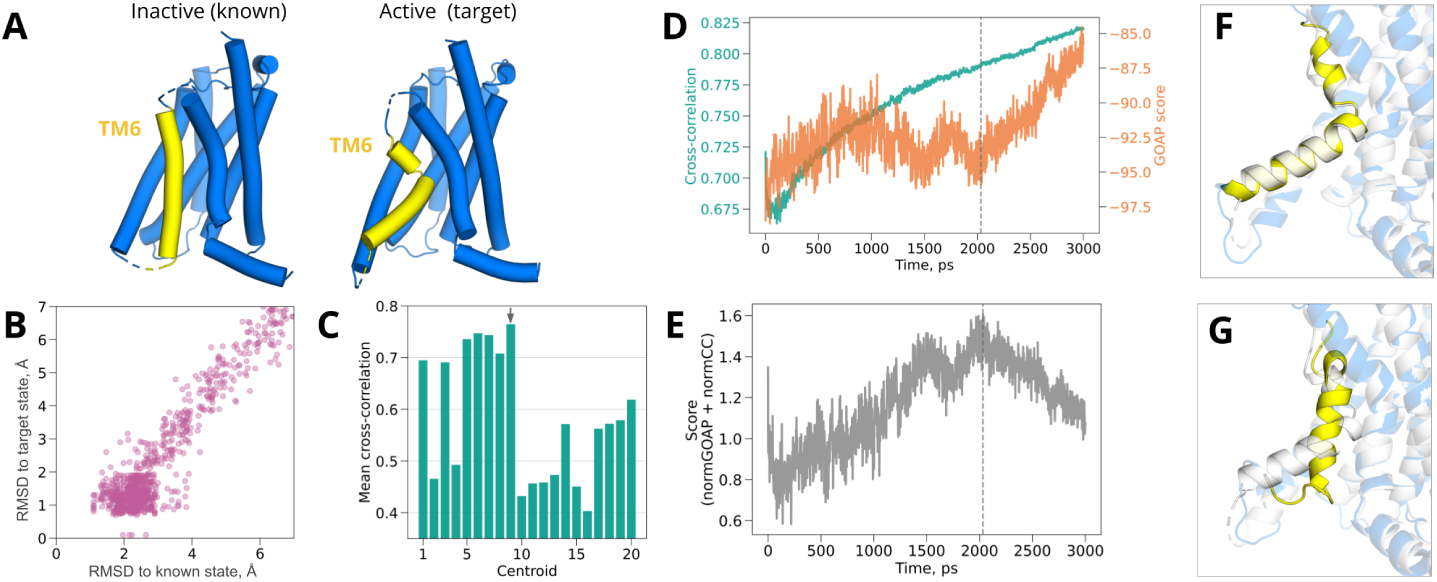
Helix bending in CLR. (A) Structures of inactive (*left*, known state, from PDB ID 7KNT [23]) and active CLR (*right*, target state, from PDB ID 6UVA [23]), with the TM6 helix — where the prominent conformational transition occurs — shown in yellow. (B) Diversity of CLR models from generated ensemble, as indicated by C*α* RMSD to target vs. known states. (C) Mean cross-correlation over density-guided simulations for each cluster centroid, with the best-fit centroid indicated by a gray arrow. (D) Time-dependent cross-correlations (green) and GOAP scores (orange) during density fitting for the best-fit cluster centroid shown in panel *C*. (E) Time-dependent compound score, combining cross-correlation and GOAP, during density fitting for the best-fit cluster centroid. In panels *D* and *E*, dashed line indicates the frame with the best compound score, selected as the final model. (F) Overlay of the target structure (white) with the model fitted from our ensemble (colored). The bulk of the protein is transparent, with the fitted model in blue; the transitional TM6 region is opaque, with the fitted model in yellow. (G) Overlay of the target structure (white) with the model fitted by standard protocol starting from the known experimental structure (colored). Same coloring scheme is used as in panel *F*.

The bulk of our generated ensemble fell within the cutoff for good quality geometry (Fig. S1). Moreover, most models deviated 1-7 Å (C*α* root-mean-square deviation, RMSD) from either the known or the target structure (PDB ID 6UVA)[20], indicating that the dataset was reasonably diverse but included conformations potentially suitable for refinement in either state (Fig. 4B). Note that deviation from the target state was monitored as an assessment of the ensemble quality, but it is not used for the refinement. After filtering by GOAP score and *k* -means clustering in MDAnalysis, several (12/20) centroids produced density-guided simulations with mean cross-correlations over 0.5 (Fig. 4C); a frame within 2.1 ns in the best-fit trajectory had the highest compound score (Fig. 4D-E) and was chosen as a final model.

We then compared to standard flexible fitting, starting from the known state for CLR in the inactivated state. The final model was within 1.3 Å C*α* RMSD from the activate experimental structure (target state) (Table 2, Fig. 3A), and included the active-state kink (Fig. 4G), such that even local deviation in TM6 was limited to 1.6 Å (Fig. S2). In contrast, the best model from standard density-guided simulations of the inactive structure (known state) deviated more than 2.3 Å from the target, both globally and locally (Table 2, Fig. 3A). Compared to standard known-state fitting, our approach also produced modestly better model quality (−96 vs. -91 normalized GOAP, 0.2 vs. 2.4 Clash, 1.4 vs. 1.9 MolProbity scores) and fit to the target density (0.79 vs. 0.75 cross-correlation) (Table 2).

**Table 2.**
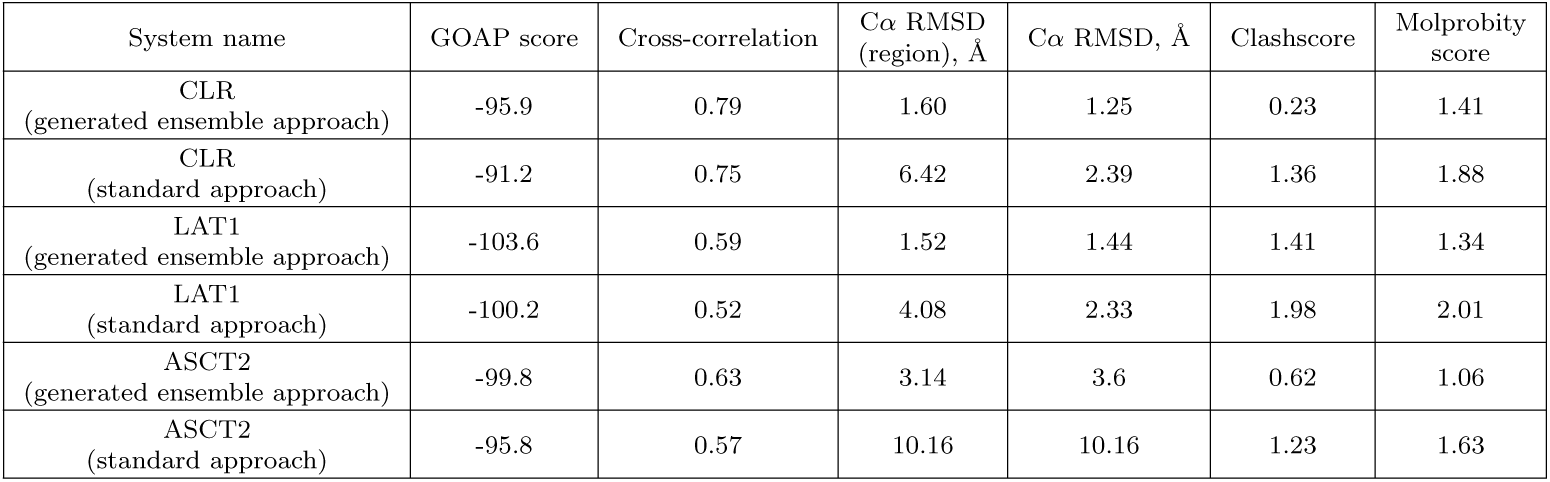
Metrics evaluating structure refinement quality.

Visual inspection of fitted models indicated that the relative success of the generative modeling approach was largely attributable to TM6. The best-fit centroid model from our generated ensemble, as well as the final model from subsequent density-fitting, included the TM6 kink associated with the active target state (Fig. 4G, Fig. S3A-B). Local deviation in TM6 was limited to 1.6 Å relative to the target structure (Fig. S2), and key side chains at the intracellular interface were correctly oriented, despite the absence of the G*α*s protein binding partner during fitting (Fig. S3A,C). Conversely, TM6 remained largely straight after density-guided simulations of the inactive known structure (Fig. 4F), such that local deviation from the target structure in this region was greater than 6 Å (Table 2 and Fig. S2). Attempted fitting of straight TM6 to the density resulted in partial unwinding of the helix N-terminus, positioning different residues at the G-protein interface than in the target structure (Fig. S3B,D). Thus, generated ensemble models offer distinct advantages in the case of a local helix transition between functional states of CLR.

### Test case 2: multiple helix rearrangements in a 12-TM transporter

To test our approach in context of a larger conformational transition, we next applied it to LAT1, the light-chain component of a heteromeric amino-acid transporter. Together with its heavy-chain binding partner 4F2hc, LAT1 carries hormones and drugs across the blood-brain barrier, and is overexpressed in several cancers [29, 30]. A member of solute carrier family 7 (also designated SLC7A5), LAT1 has a 12-TM LeuT fold, and cycles between various inward, occluded, and outward states in a rocking bundle mechanism to transport solutes across the plasma membrane [31]. We used a structure of LAT1 in the inward-open configuration, extracted from the heterodimeric complex (PDB ID 6IRS) [24], as the known state (Table 1). As a target, we used a cryo-EM density (EMD 30841) thought to represent an intermediate between outward-occluded and outward-open states, taken from a complex with the inhibitor 3,5-diiodo-L-tyrosine (diiodo-Tyr) [21]. The principal distinction between the known and target states involved rearrangement of the kinked helices TM1 and TM6 (residues 47-80 and 240-265 respectively in the target structure) around the diiodo-Tyr binding site (Fig. 5A, Fig. S4). The protein density was reported at lower overall resolution than that of CLR (3.4 Å vs. 2.3 Å), offering an additional test case for lower-quality data. We still added 1 Å Gaussian blur to the density.

**Fig. 5.**
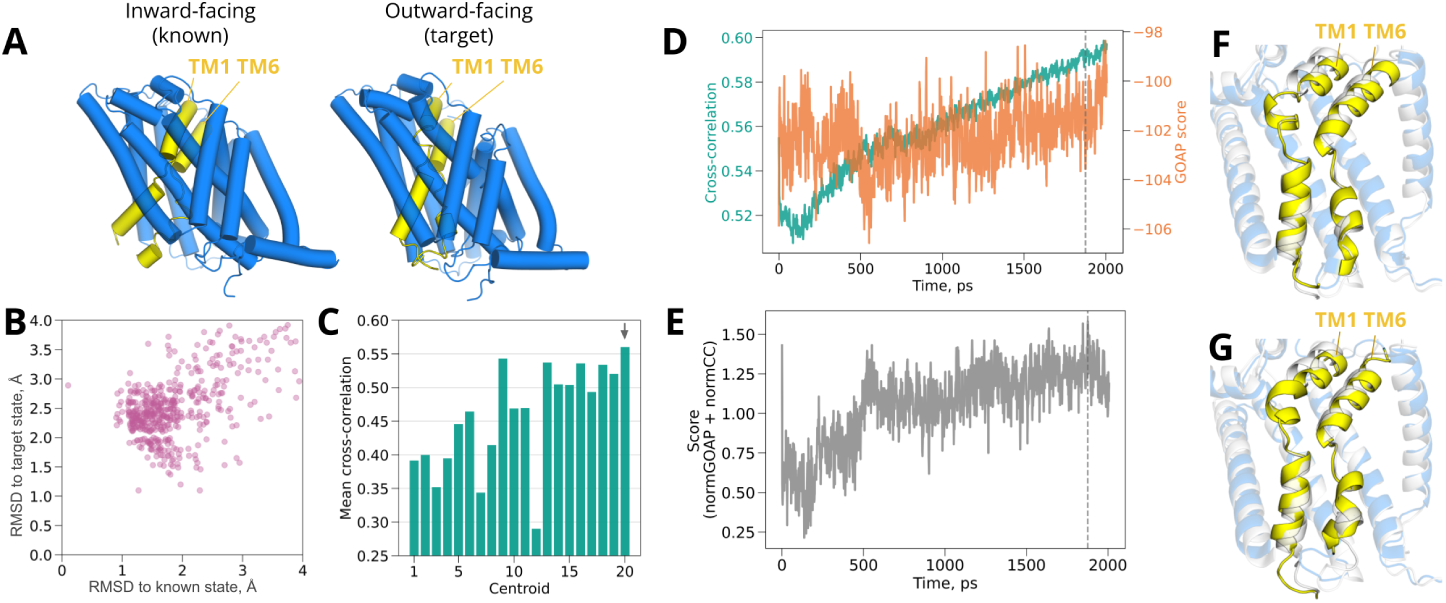
Rearrangement of helices in LAT1. (A) Structure of inward-open (*left*, known state, from PDB ID 6IRS [24]) and inhibited LAT1 (*right*, target state, from PDB ID 7DSQ [21]), with the TM1 and TM6 helices — where the prominent conformational transition occurs — shown in yellow. (B) Diversity of LAT1 models from generated ensemble, as indicated by C*α* RMSD to target vs. known states. (C) Mean cross-correlation over density-guided simulations for each cluster centroid, with the best-fit centroid indicated by a gray arrow. (D) Time-dependent cross-correlations (green) and GOAP scores (orange) during density fitting for the best-fit cluster centroid shown in panel *C*. (E) Time-dependent compound score, combining cross-correlation and GOAP, during density fitting for the best-fit cluster centroid. In panels *D* and *E*, dashed line indicates the frame with the best compound score, selected as the final model. (F) Overlay of the target structure (white) with the model fitted from our generated ensemble approach (colored). (G) Overlay of the target structure (white) with the model fitted by standard density-guided simulations of the known structure (colored). In panels *F* and *G*, the bulk of the protein is transparent, with the fitted model in blue; the transitional TM1/TM6 region is opaque, with the fitted model in yellow.

The results indicated that our LAT1 target density was more challenging than CLR, but could be accurately modeled. Our generated ensemble for LAT1 included more models that exceeded the cutoff for reasonable quality geometry (Fig. S1); on the other hand, LAT1 models deviated less than 4 Å (C*α* RMSD) from either the known or target (PDB ID 7DSQ [21]) structures, indicating the presence of conformations similar to both states (Fig. 5B). Cluster centroids were successfully fitted using density-guided MD simulations, with 8/20 centroids showing over 0.5 mean cross-correlation (Fig. 5C); a final model was selected within 1.8 ns in the best-fit simulation on the basis of compound score (Fig. 5D-E).

As in the case of CLR, our approach resulted in better fit and geometry metrics than standard density fitting of the known LAT1 structure. The final model based on our generated ensemble was within 1.5 Å C*α* RMSD from the inhibited target structure (PDB ID 7DSQ [24]), while the best model based on fitting the inward-open known structure deviated more than 2.3 Å (Table 2, Fig. 3B). Our pipeline model also deviated by only 1.5 Å in the TM1/TM6 region, and was compatible with binding diiodo-Tyr, despite the absence of this inhibitor model and its density during fitting (Table 2, Fig. S4). Conversely, the known-state fitted model deviated over 4.0 Å (Table 2) in the TM1/TM6 region, and oriented a Tyr residue to clash with the anticipated inhibitor pose. Compared to standard known-state fitting, our generative ensemble setup produced modestly better model quality (−104 vs. −100 normalized GOAP, 1.4 vs. 2.0 Clash, 1.3 vs. 2.0 MolProbity scores) and fit to the target density (0.59 vs. 0.52 cross-correlation). Thus, our approach facilitated modeling in the context of a multi-helix conformational transition, including a ligand binding site in a functional intermediate.

### Test case 3: domain rearrangement in an 8-TM transporter

To test the applicability of our approach to a more distributed transition between known and target states, we investigated ASCT2, a homotrimeric member of solute carrier family 1 (SLC1A5) with an 8-TM GltPh fold [32]. This transporter is widely expressed in human tissues, and upregulated in numerous cancers, making it an important target for drug development as well as biophysical characterization [33]. ASCT2 is structurally and mechanistically distinct from LAT1, consisting of a scaffold domain comprising helices TM1-2 and TM4-5, and a transport domain comprising TM3 and TM6-8 along with two interhelical hairpins [34]. Transport occurs by a one-gate elevator mechanism, in which the transport domain moves between inward and outward states relative to the scaffold [35]. We used a structure of an ASCT2 monomer in an inward-open configuration, extracted from the trimeric complex (PDB ID 6RVX) [25], as the known state (Table 1). As a target, we used a cryo-EM density (EMD 12142) determined to similar resolution as LAT1 (3.4 Å) in an outward-open configuration, taken from the homotrimeric complex in the presence of the inhibitor 4-(4-phenylphenyl)carbonyloxypyrrolidine-2-carboxylic acid (Lc-BPE) [22]. Conformational cycling in this system involved evident rearrangements among all helices and hairpins; accordingly, we considered the entire monomer as a transition region (Fig. 6A).

**Fig. 6.**
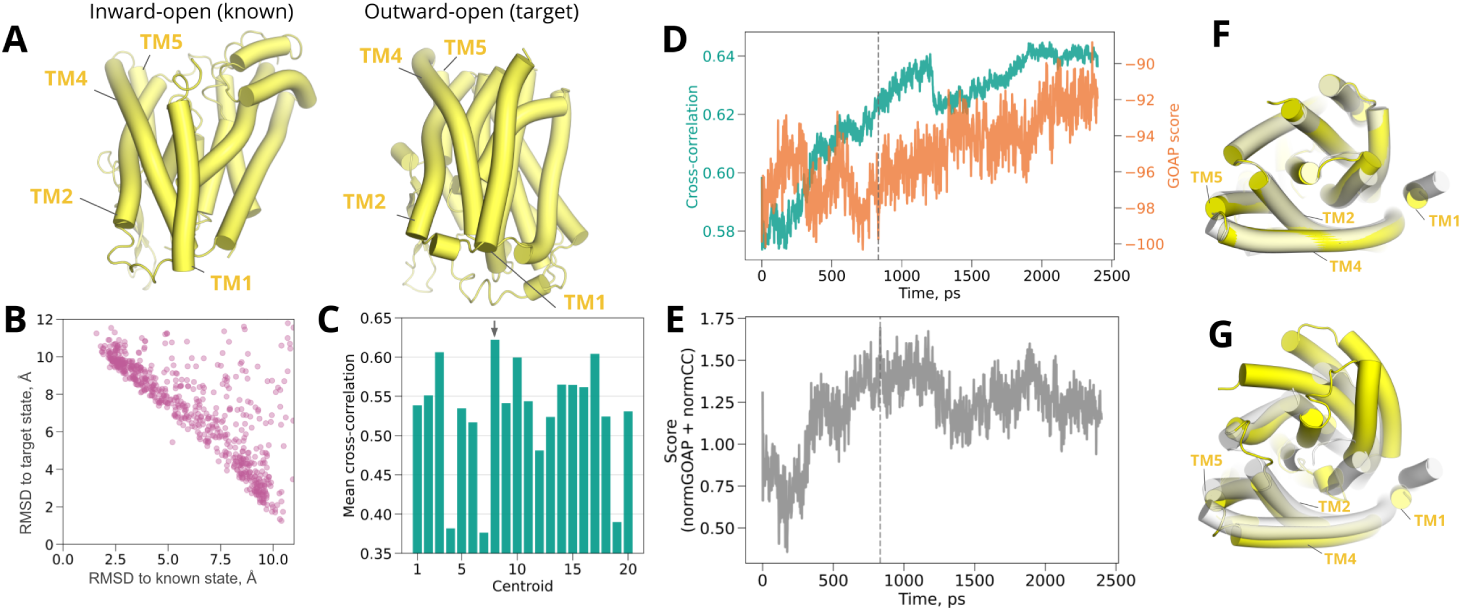
Domain rearrangement in ASCT2. (A) Structures of inward-open (*left*, known state, from PDB ID 6RVX) [25] and outward-open ASCT2 (*right*, target state, from PDB ID 7BCQ [22]). (B) Diversity of ASCT2 models from generated ensemble, as indicated by C*α* RMSD to known vs. target states. (C) Mean cross-correlation over density-guided simulations for each cluster centroid, with the best-fit centroid indicated by a gray arrow. (D) Time-dependent cross-correlations (green) and GOAP scores (orange) during density fitting for the best-fit cluster centroid shown in panel *C*. (E) Time-dependent compound score, combining cross-correlation and GOAP, during density fitting for the best-fit cluster centroid. In panels *D* and *E*, dashed line indicates the frame with the best compound score, selected as the final model. (F) Overlay of the target structure (white) with the model fitted from our generated ensemble approach (yellow). (G) Overlay of the target structure (white) with the model fitted by standard density-guided simulations of the known structure (yellow). In panels *F* and *G*, helices contributing to the scaffold domain are labeled; remaining helices constitute the transport domain.

Despite its relatively distributed transition between known and target states, our approach was similarly successful in modeling ASCT2 as previous test cases. Our generated ensemble included an intermediate fraction with low quality geometry (Fig. S1); however, models within 2 Å C*α* RMSD of the known structure deviated at least 10 Å from the target (PDB ID 7BCQ [22]) and vice versa, consistent with substantial rearrangements observed between the two states (Fig. 6B). Most (16/20) cluster centroids could be fit with mean cross-correlation to the target map over 0.5 (Fig. 6C), with the final model selected within 0.8 ns in the best-fit simulation (Fig. 6D-E).

Fitting of ASCT2 was substantially more accurate using our ensemble approach than the known structure (Fig. 3C). The final model based on our generated ensemble was within 3.2 Å C*α* RMSD from the outward-open target structure, while the best model based on fitting the inward-open known structure deviated more than 10.1 Å (Table 2). When aligned on the scaffold domain (TM1-2, TM4-5), all transmembrane helices in our final model were largely superimposable with the target structure (Fig. 6F); for the fitted known state, rearrangements were evident in helices comprising the transport domain (Fig. 6G). As in previous cases, our approach also produced modestly better model quality (−100 vs. −96 normalized GOAP, 0.6 vs. 1.2 Clash, 1.1 vs. 1.6 MolProbity scores) and fit to the target density (0.63 vs. 0.57 cross-correlation). Thus, generated ensemble model fitting appeared particularly useful in the context of a conformational transition across multiple transmembrane helices.

## Discussion

In this study, we propose and implement an approach that leverages generative protein models along with density-guided MD simulations for automated atomistic modeling into cryo-EM maps, applicable among other things to multi-state proteins. Automated structure determination often relies on rigid or flexible fitting of known structures; however, even flexible fitting may fail to accurately refine a complex target for which the initial model substantially differs. To illustrate this concern, our test cases include three pharmacologically relevant membrane proteins for which cryo-EM structures are available in at least two conformations. Activation by G protein binding to CLR involves bending of a key transmembrane helix; inhibition by diiodo-Tyr locks LAT1 in an intermediate, with two transmembrane helices rearranged relative to the inward-open state; and conformational cycling between inward-open and outward-open states involves distributed remodeling across ASCT2, particularly in the transport relative to scaffold domains.

In our test cases, standard density-guided MD simulations based on one (known) structure result in models that deviate from the target structure by 4.1-10.2 Å. In contrast, flexible fitting using many alternative full-length models generated in AlphaFold2 as starting points recapitulated target structures within 1.6-3.2 Å. Notably, this generated-ensemble approach removes the need to modify initial models for density fitting, for example by building atoms or residues unresolved in a previous experimental structure. Although this approach may prove less successful for systems with limited structural data, all our target systems were excluded from the AlphaFold2 training set, and in fact have been used in previous work to validate the applicability of this tool to new systems [18]. In cases for which restricted homology or intrinsic disorder prevents approximation of the target conformation [36], multi-step approaches using progressive resolution [37] or enhanced sampling [10, 11] may prove valuable.

Our approach further leverages stochastic subsampling of MSA depth to generate more diverse initial models, allowing us to sample conformations suitable for automated fitting to the target state. Some de-novo methods of cryo-EM structure determination incorporate prediction tools such as AlphaFold or RoseTTAFold to generate protein or fragment models[38]; such functionality is available in several protocols and tools [39–49]. However, even when using generative models, predicting divergent or rare protein conformations may be challenging [50]. By instead generating and clustering a substantial and diverse ensemble of full-length models, it is sufficient that *one* of the cluster centroids is sufficiently close to the density to enable accurate refinement. Thus, this approach can be particularly valuable for proteins with multiple metastable states or otherwise flexible features. Whereas MSA subsampling proved effective in our test cases, other recent approaches to generating diverse structural ensembles in AlphaFold2 [51, 52] or AlphaFold3 [53] may offer further advantages. In particular, sequence clustering or alignment alterations may improve sampling or allow tuning to specific targets or transitions. Although the alphafold2 conformations package used here does not support oligomeric proteins [18], tools such as AlphaFold2-multimer may enable the application to larger, more complex systems [54]. Alternative approaches to conformational sampling include extensive MD simulations, but may be substantially more resource intensive than the generative modeling approach taken here.

Integrating flexible fitting with ensemble generation offers further potential advantages [55, 56]. Whereas classical de-novo methods [5–7] work reliably with high-resolution cryo-EM data, they may be less applicable to medium- and low-resolution maps. In such cases, density-guided MD simulations may be particularly valuable in integrating physics-based force fields with cryo-EM data. In fact, higher resolution densities are often blurred for the purpose of flexible fitting, reducing barriers to translocate model regions from one density area to another [10]. Target densities tested in this work were resolved to 2.3–3.4 Å overall, and each were subjected to an additional 1 Å Gaussian blur prior to fitting. Furthermore, reducing our ensemble to cluster centroids allowed us to run only 20 density-guided simulations for each system, while representing the diversity of 1250 models. Even where the centroid may not have represented the best model, application of the MD force field during flexible fitting evidently enabled us to optimize geometry alongside map correlation. We were able to generate and fit models for each system with relatively high accuracy despite the absence of secondary structure restraints or membrane mimetics, further streamlining the protocol. Manual verification of the resulting models might be appropriate prior to final deposition. Although our test systems were extracted from larger complexes, flexible fitting in the absence of such partners nonetheless recapitulated key interfaces with accessory G proteins (for CLR) or inhibitors (for LAT1). Thus, our approach offers a straightforward integration of the strengths of both model generation and flexible fitting, applicable even to membrane-protein systems representing a range of data quality and binding partners.

## Methods

### Test systems

Three different systems, each including cryo-EM structures in at least two different conformations, were selected as known and target states for pipeline validation: inactive and active conformations of CLR transmembrane domain [20, 23], inward-open and inhibited conformations of LAT1 [21, 24], and inward-open and outward-open conformations of ASCT2 [22, 25]. All target structures and densities were released after the training dataset for AlphaFold v.2.0.1.

### Ensemble generation

AlphaFold2 models were generated with the alphafold conformations package [18]. Before ensemble generation, we assessed per-residue prediction quality for each protein sequence using the AlphaFold Protein Structure Database [57]. Terminal residues with low-confidence predictions (pLDDT scores below 70) were trimmed from the sequence to improve further modeling reliability and exclude regions unlikely to correspond to well defined densities. MSAs were obtained from the MMseqs2 database [58]. The MSA depth was subsampled according to previously proposed methods [18], using a single recycle and no energy minimization. MSA depths in the range of 5-30 were used, with 50 models generated for each depth, giving 1250 models for each target.

### Model clustering and rigid alignment

After model generation, we filtered out models that were completely or partially mis-folded on the basis of GOAP score [59] normalized over protein sequence length. GOAP source code was downloaded at https://sites.gatech.edu/cssb/goap/. We dis-carded models with normalized scores above -100, a threshold value selected based on GOAP distributions (Fig. S1) and visual evaluation. The remaining models were aligned to the structure assigned as the known state using PyMOL [60], and clustered using *k* -means clustering in the MDAnalysis Python package[61] with target n clusters=20 and other parameters as default. Rigid alignment of cluster centroids to the cryo-EM density assigned as the target state was performed using the rigidbodyfit 1.2.1 package available at https://gitlab.com/cblau/rigidbodyfit.

### Density-guided simulations

Initial preparation of the target density was performed in UCSF Chimera [62]. Preparation included segmentation of the original density map and manual picking of the segments corresponding to the protein of interest. After merging selected regions, Gaussian filtering was performed with standard deviation of the 3D-Gaussian function of 1 Å. Missing loops in the known experimental states of CLR and LAT1 were built using MODELLER [63] in UCSF ChimeraX [64]. System preparation, equilibration and density-guided simulations were performed in GROMACS 2023.1 [65]. TIP3 water, the neutralising amount of NaCl, and the CHARMM36 force field [66] were used for all simulations. It has been shown that embedding in lipids or detergents does not substantially improve density-guided simulations of a membrane protein [67], so for simplicity they were not included. Bonded and short-range nonbonded interactions were calculated every 2 fs, and periodic boundary conditions were employed in all three dimensions. The particle mesh Ewald (PME) method [68] was used to calculate long-range electrostatic interactions. A force-based smoothing function was employed for pairwise nonbonded interactions at 1 nm with a cutoff of 1.2 nm. The temperature was maintained at 300K with the velocity-rescaling thermostat [69]. The pressure was kept at 1 bar with the c-rescale barostat [70]. Forces from density-guided simulations were applied every *N* = 2 steps. The scaling factor for density-guided simulation forces of 10^3^ kJ/mol was combined with adaptive force scaling. All simulation setup parameters have been made available [71].

### Analysis

For each target, we identified the simulation with the highest mean cross-correlation to the target density, then selected the frame with the highest compound score as a final model. Standard cross-correlation calculations were performed using a modified GROMACS version, available at https://gitlab.com/gromacs/gromacs/-/tree/ ml fscavg main. The compound score was calculated by subtracting the normalized GOAP score (as described above) from the cross-correlation, enabling optimization of protein geometry as well as map fit. Phenix MolProbity clash scores and total scores were calculated for additional model quality assessment [72]. RMSD was calculated in PyMOL [60]. Python 3.9 scripts were utilised to combine pipeline steps, conduct analyses, and generate figures. Deviation in regions of local conformational change was calculated for residues 329-354 for CLR, and for residues 47-80 and 240-265 for LAT1; for ASCT2, the entire polypeptide was consider the region of change.

## Acknowledgments

Computational resources for simulations were provided by the National Academic Infrastructure for Supercomputing in Sweden (NAISS 2024/3-49). AlphaFold2 runs were performed using the computing facilities of the Berzelius through NAISS (grant no. Berzelius-2023-244).

## Funding

This work was partially funded through a Marie Sklodowska-Curie Post-doctoral Fellowship 101107036 to NH and grants from the Swedish Research Council (VR; 2019-02433, 2021-05806), the Knut and Alice Wallenberg foundation (KAW; 2023.0254) and the BioExcel-3 Centre-of-Excellence (EuroHPC Joint Undertaking; 101093290) to EL.

## Author Contributions

Conceptualization: NH, TS, RJH, EL

Data curation: TS, NH

Formal analysis: TS

Funding acquisition: NH, EL

Investigation: TS Methodology: TS, NH

Project administration: NH, RJH, EL

Resources: NH, EL

Software: TS, NH Supervision: NH, RJH, EL

Validation: TS Visualization: TS

Writing – original draft: TS

Writing – review & editing: TS, NH, RJH, EL

## Competing Interests

The authors declare that they have no competing interests.

## Data and Materials availability

All data needed to evaluate the conclusions are present in the paper or the Supplementary Information. The generated models with ChimeraX sessions, density-guided simulation input files and trajectories can be found on Zenodo: 14749350 (https://zenodo.org/records/14749350).

## Notes

https://zenodo.org/records/14749350

### Competing Interest Statement

The authors have declared no competing interest.

### Summary of Updates

The language of the manuscript has been updated to clarify some of our methodological approaches.

