## Supplementary Information for "Modeling cryo-EM structures in alternative states with generative AI and density-guided simulations"

### Refining unknown conformations in cryo-EM through ML-driven model generation and density guided simulations

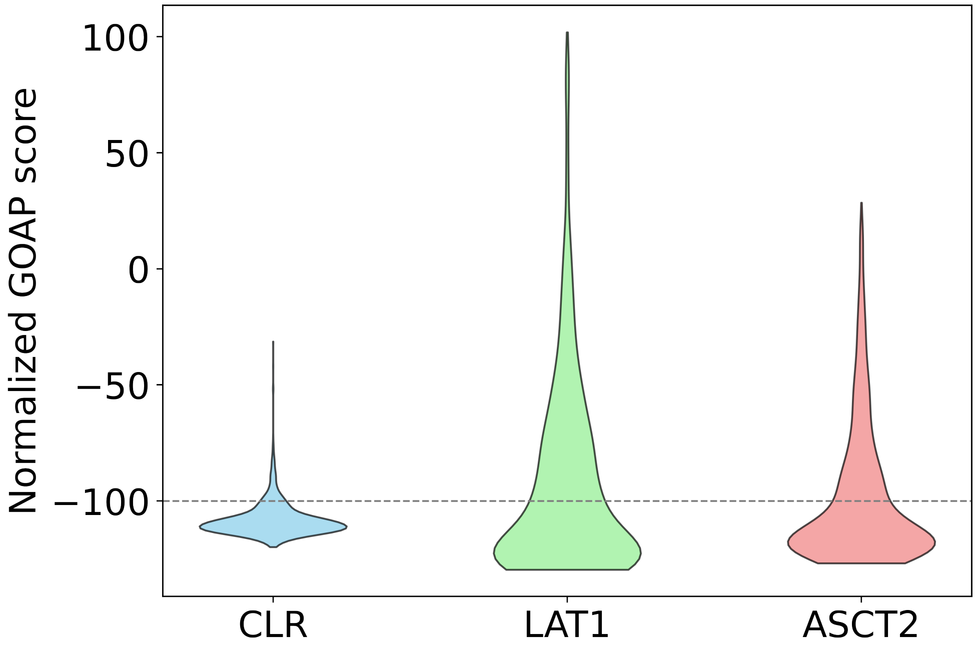


**Fig. S1 Distribution of GOAP scores normalized by protein sequence length, calculated for models produced by AlphaFold2 for each system.** The filtering threshold of -100 is depicted as a gray dashed line.

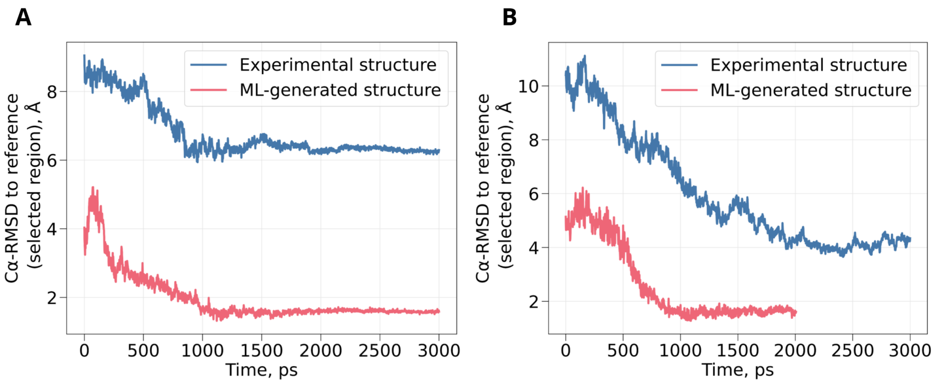


**Fig. S2 Refinement of local conformational changes in two test systems.** (A) C$\alpha$ RMSD to the target structure during density-guided simulations of the known structure (blue) or best-fit cluster centroid from our generative-AI ensemble (red) for the TM6 helix in CLR. (B) RMSD plots as in *A* for the TM1/TM6 region in LAT1.

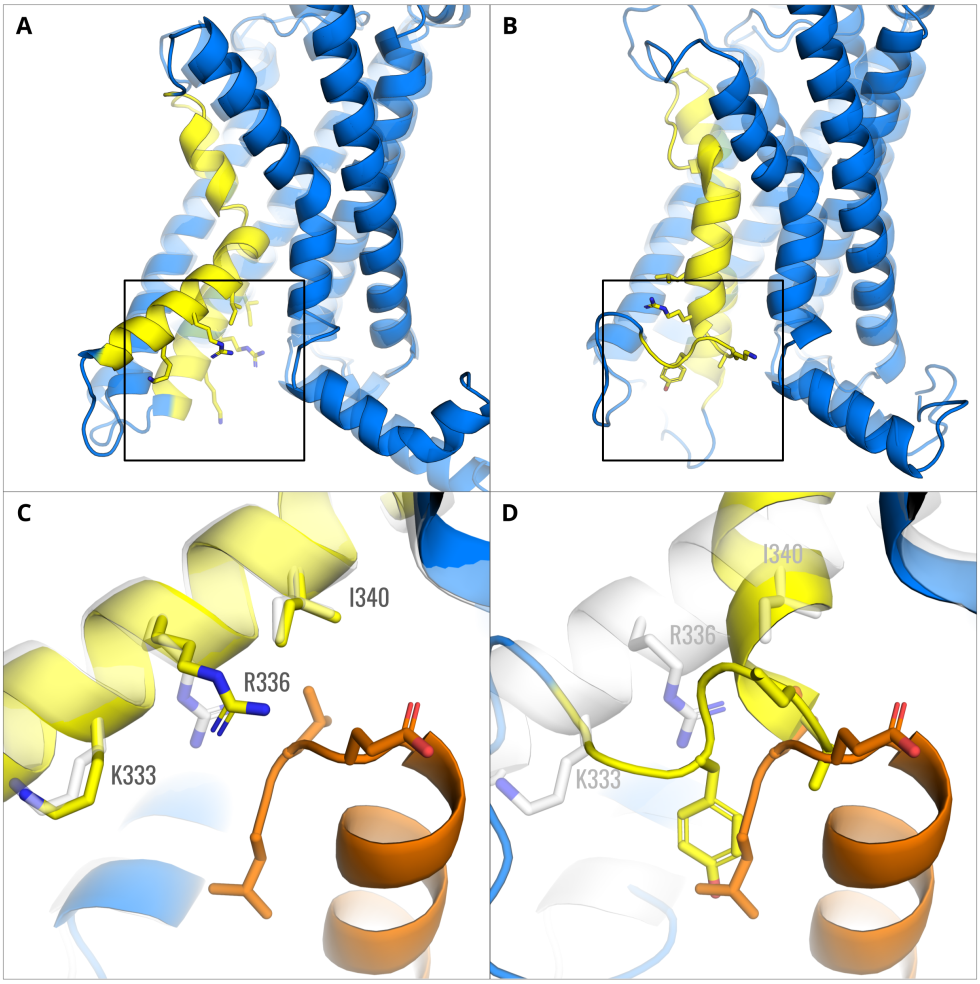


**Fig. S3 Refinement of local conformational change in CLR.** (A) Overlay of the final model from our generative-AI approach (opaque) with the corresponding initial cluster centroid (transparent). The bulk of the protein is shown as blue ribbons, with the TM6 region subject to kinking upon G-protein activation as yellow ribbons. Amino-acid residues poised to interact with the Gs-protein $\alpha$ subunit are shown as sticks, colored by heteroatom. (B) Overlay as in *A* of the model from standard known-state fitting (opaque), overlaid with the corresponding inactive structure (transparent). (C) Zoom view of the boxed region in *A*, showing the interface between TM6 and the G protein (orange). The final model from our approach (opaque, colored) is overlaid with the target structure (transparent, white). (D) Zoom view of the boxed region in *B*. The final model from the standard approach (opaque, colored) is overlaid with the target structure (transparent, white).

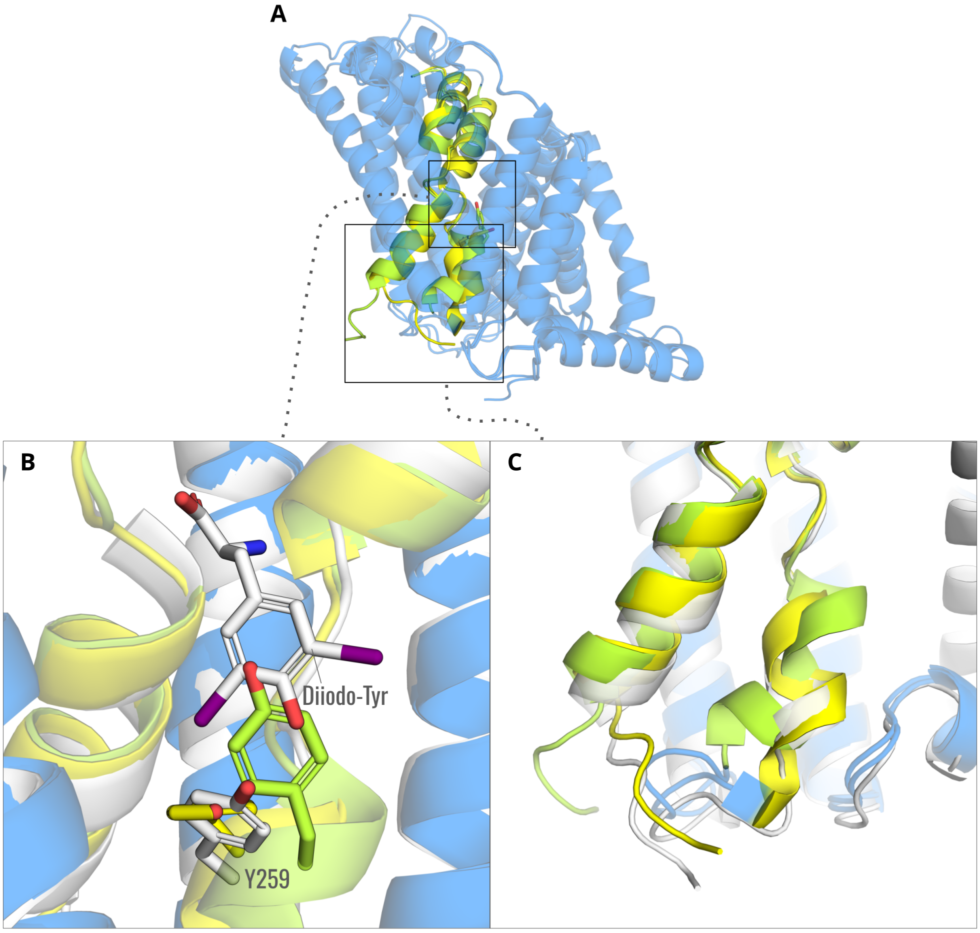


**Fig. S4 Refinement of local conformational change in LAT1.** (A) Overlay of final models from generative-AI and standard pipelines. The bulk of both proteins is shown as blue ribbons, with the TM1/TM6 region subject to rearrangement in the inhibited state colored differently for fitting based on the best-fit cluster centroid (yellow) and fitting based on the known structure (green). Residue Y259 is shown as sticks, colored by heteroatom. (B) Zoom view of the upper boxed region in *A*, including the target structure (white ribbons) at the interface between Y259 and diiodo-Tyr (white sticks). Ribbons and sticks for the fitted models are otherwise colored as in *A*, showing prospective clash with the inhibitor in the standard approach. (C) Zoom view of the lower boxed region in *A*, showing the intracellular end of the TM1/TM6 region. Local rearrangements are relatively poorly fit to the target structure (white) by the standard (green) versus generative-AI (yellow) pipelines.
